# VRmol: an Integrative Cloud-Based Virtual Reality System to Explore Macromolecular Structure

**DOI:** 10.1101/589366

**Authors:** Kui Xu, Nan Liu, Jingle Xu, Chunlong Guo, Lingyun Zhao, Hong-Wei Wang, Qiangfeng Cliff Zhang

**Author notes:** These authors contributed equally to this work (Kui Xu, Nan Liu).

## Abstract

Past decades have seen much effort devoted to the tool development for biomacromolecule structural visualization and analysis. In recent years, we have witnessed tremendous advancement in computational technologies, especially in graphics and internet. Here we present an integrative cloud-based, virtual reality (VR) system -- VRmol that exploits these technology advances to visualize and study macromolecule structures in various forms. The immersive environment can improve user experiences in visualization, calculation, and editing of complex structures with more intuitive perception. VRmol is also a system with cloud-based capability to directly access disease-related databases, and couples the structure with genomic variation as well corresponding drugs information in a visual interface, thus serving as an integrative platform to aid structure-based translational research and drug design. VRmol is freely available at https://VRmol.net.

## INTRODUCTION

Graphical visualization and analysis of molecular structures can yield insights into the mechanisms of actions of macromolecular machines. GRASP [1], RasMol [2], PyMol [3] and UCSF Chimera [4] are among widely used software tools for molecular visualization with comprehensive functions for structural analysis. However, tools designed for desktop systems are often tedious to install, configure and use, and are inconvenient to share real-time graphic scenes with collaborators using different computers or physically away from each other. Although some web-based applications like JMol [5] and AstexViewer [6] are straightforward, they only support limited types of structural data and cannot perform complex tasks of structural analysis. In particular, recent progresses in structural determination by cryo-EM demands novel web-based tools that efficiently support cryo-EM maps and structural models of large complexes [7, 8]. More importantly, new types of genomics and translational information are accumulating online as research enters the post-human genome project era [9]. There is a pressing need to develop an integrative platform that can combine these big data to facilitate the investigation of biological structural systems. With the recent developments in computer graphics and internet technologies, it is now possible to fill these gaps by developing powerful web-based tools that offer both high quality graphical visualization and complex structural analysis.

Virtual reality (VR) is a new type of computer graphics technology that creates a virtual world, enabling people to interact with simulated objects beyond reality [10, 11]. Areas such as engineering, education, arts, entertainment, and biomedical research have started to embrace VR for better visualization, modeling, and simulation [12, 13]. Recently, VR systems have been used to facilitate molecular simulation by creating an interactive way to help determine the simulation strategy [14, 15]. AltPDB protein-viewer provides an immersive VR environment for viewing and for collaborative studies on complex biological structures. However, structural analysis by AltPDB protein-viewer is complicated due to the need to generate molecular models beforehand by third-party tools. UCSF ChimeraX [16, 17] also provides certain simple VR interactive functions for structure visualization, but it is mainly designed for a native desktop environment and time-consuming to learn. In addition, some favorable structural analysis options, like molecules docking and scene sharing, are inaccessible in the VR environment of ChimeraX.

In parallel to the advances in graphics, internet is another fast-developing computational area with many inspiring technology innovations. Web-based tools with flexible and simple interfaces to study macromolecular structures have emerged, such as AstexViewer [6], Jmol [5], NGL Viewer [18], 3Dmol.js [19], LiteMol [20] and Web3DMol [21]. With the help of the Web Graphics Library (WebGL), it is now possible to create web-based VR experiences that can be directly accessed with common web browsers. A recently developed web tool, Autodesk Molecular Viewer [22], implemented a VR environment with Cardboard VR device based on browsers on smartphones, like Chrome or Safari. However, some basic functions of structural analysis are absent, such as molecular surface representation and EM density map visualization, not to mention more advanced analytical tasks, such as structural conservation study and molecular structure fragmentation.

Importantly, genomics technologies and translational researches have generated a huge amount of data that could inform disease studies and drug design. Databases like OMIM [23], HGMD [24], and ClinVar [25] have been created to curate and record disease-or phenotype-associated genomic mutations. Systematic studies of disease and cancer genomes have produced an explosive growth of human genomic variation data, exemplified by The Cancer Genome Atlas (TCGA) [26], the Cancer Cell Line Encyclopedia (CCLE) [27], and the Exome Aggregation Consortium (ExAC) [28]. Mapping and studying the mutations and variations in a structural context can shed light on the molecular mechanisms of related diseases [29]. Integrative analysis with corresponding potential drug molecules from databases, like the DrugBank [30] and ChEMBL [31], may yield insights for drug action mechanisms and efficacy, and further optimization. However, to the best of our knowledge, no existing molecular viewing tool integrates structural analysis with these genomic variation and drug molecule databases.

Addressing these gaps, we developed a novel web-based framework, VRmol, for molecular structure visualization and integrative analysis. Moreover, VRmol connects to multi-source diseases and drugs related database from the cloud in a fully immersive VR environment, providing an integrated platform for translational study. Users can access VRmol at https://VRmol.net through a WebVR enabled web browser, such as Microsoft Edge and Firefox Nightly.

## MATERIALS AND METHODS

### System Design and Implementation

The VRmol system consists of three modules: (1) a resource module to retrieve and parse different types of structural and translational data from the cloud by connecting to online databases including the protein data bank (PDB) [32], electron microscopy data bank (EMDB) [33], disease-related genomic mutation and variations data (like HGMD, ClinVar and TCGA), and drug databases (like DrugBank and ChEMBL); (2) a structure modeling module to generate, process and render 3D structural models in VR environment implemented by native JavaScript with WebGL, WebVR and Three.js (Supplementary Table S1); (3) an interaction module to analyze, manipulate and interact with structural models in an immersive VR environment using different VR devices such as HTC Vive, Oculus Rift, and Microsoft Mix Reality in compatible web browsers (Figure 1, Supplementary Table S2).

**Figure 1.**
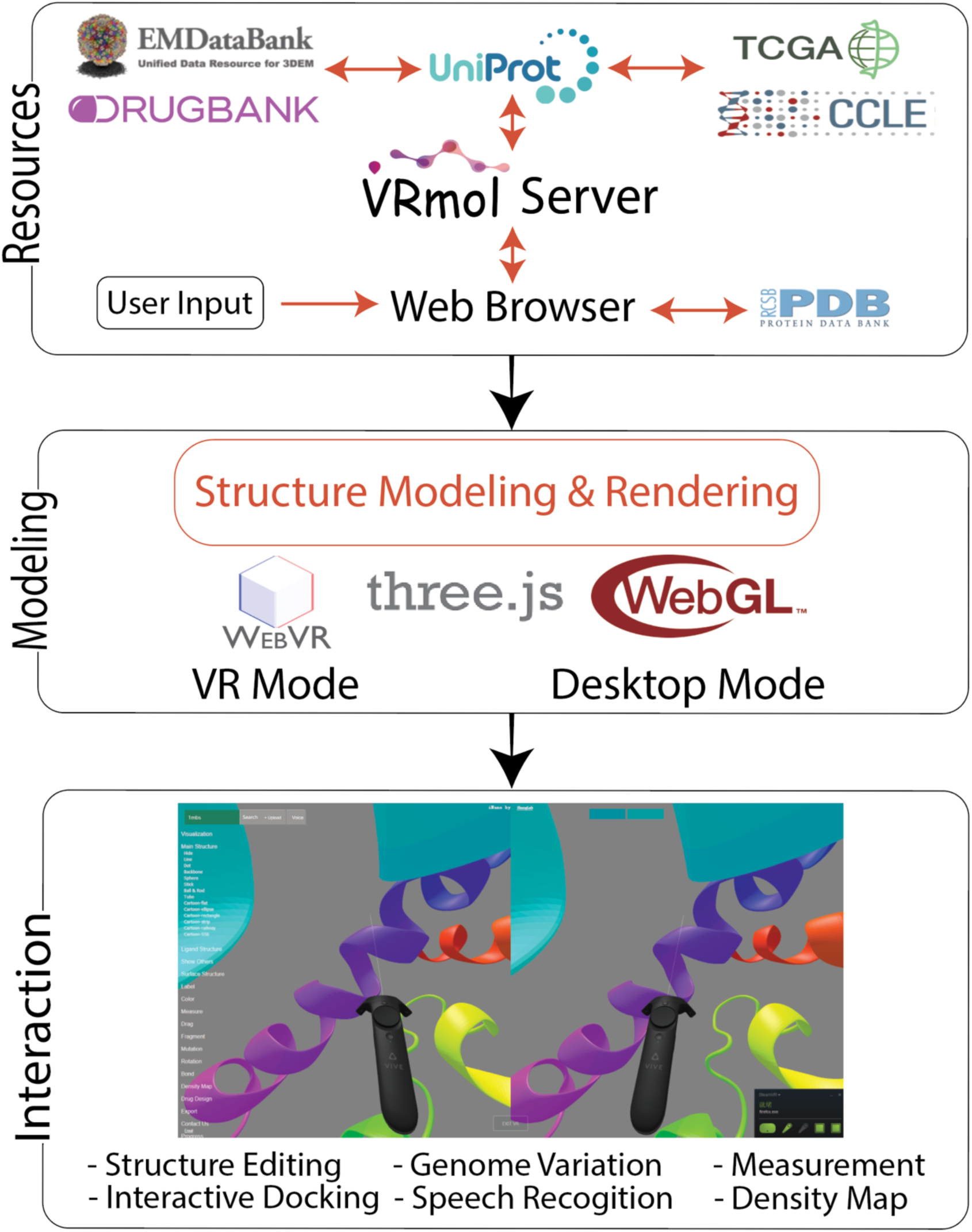
System design of VRmol. Resource module: VRmol integrates multiple structural and translational databases including PDB, EMDataBank, Uniprot, TCGA, CCLE, DrugBank etc. Modeling module: Geometry graphics of structural models are generated in a 3D virtual scene by applying Three.js library with WebGL and WebVR. Interaction module: VRmol implements multiple functions to view, analyze and edit a structure in the VR mode.

### Molecular Structural Data Parsing and Modeling

The parsing of the structure file in VRmol is format-dependent. For structural files in the PDB format (http://www.wwpdb.org/documentation/file-format-content/format33/v3.3.html),VRmol obtains the meta-information as well as the 3D coordinates by following a previously published method in Web3DMol [21], and assembles the structure geometry based on the each atom information. Briefly, the main chains are built by using the Cubic Hermite spline to interpolate 3D coordinates of the main-chain atoms, and the secondary structures in ribbon styles are achieved by extruding a 2D shape along the main chain, using the *THREE.ExtrudeGeometry* function in Three.js, a WebGL-based library. VRmol follows Web3DMol to create the tortuosity frames of an alpha-helix and a beta-sheet by computing the tangents, normals and binormals on the spline. VRmol uses the matching cube algorithm for surface modeling, including the Van der Waals surface, the solvent accessible surface, the solvent excluded surface, and the molecular surface, as described in GLmol ((https://github.com/biochem-fan/GLmol). The color schema is consistent with that in Pymol [3] and Web3DMol. For coloring by sequence conservation, VRmol uploads the structure file onto the ConSurf server [34] to compute a conservation score for each residue, based on which the residues are variedly colored. Notably, to improve the modeling efficiency of macromolecules with a large number of atoms, VRmol defines a sphere centered at the camera position. The part of the molecular model inside the sphere is rendered in high resolution (Each atom is modeled with 760 triangular plates), while the outside part is rendered in low-resolution (Each atom is modeled with only 112 triangular plates) or hidden.

For electron density maps in the MRC/CCP4 2014 format (http://www.ccpem.ac.uk/mrc_format/mrc2014.php), VRmol firstly extracts the header information and a 3D array (*M*) with density value from the input structural file, and generates three representation styles: surface (including meshed surface), dot and slice. The surface representation style is implemented by applying matching cube algorithm (as described above), and the dot style is calculated based on the voxels with density values larger than a specific threshold *t* (*t*= mean(*M*) by default). The slice style is drawn as a heat map to represent the selected plane in the 3D array. For large maps, VRmol will perform an adaptive down-sampling before structural modeling to downscale the map for efficiency.

### VR Rendering

Once the structural model is created, VRmol renders the appearances of the model, such as colors, texture, interaction, animation, and lighting environment, with the “stereo rendering” method [35] implemented with Three.js. Briefly, VRmol used the function *THREE.VRDisplay* to create VR scenes, and *THREE.VREffect* to render molecular structure models. Again, for efficnecy, VRmol also implements a spherical view in the VR scene.

### Fragmentation

VRmol includes an important function, *Fragment*, which allows to highlight selected fragments in the input structure, named as fragments of interest (FoI), by individually presenting them. Because VRmol models structures by individual atoms, FoI could be defined as any sets of residues, e.g, those of a specific class, a type of secondary structures, a region, or a chain. Every residue in the molecule has three attributes: *representation, color schema* and *selection*. The attribute *selection* of all the amino acids in a FoI are set to ‘true’, while others are ‘false’. Notably, multiple FoIs could be simultaneously selected for visualization, analyzing and modification. The edited model can be exported as a new PDB file.

### Mapping Genomic Variations to Protein Structures

Genomic mutation files (*.maf format) are downloaded from TCGA/CCLE/ExAC, annotated by oncotator (http://portals.broadinstitute.org/oncotator/), and imported into the mutation database on the VRmol server. For a query structure with a PDB code, VRmol server will map the code to identify the corresponding mutation records in the mutation database. The resulting mutation sites are then labeled with twinkling balls on the query structures, along with the related information including the disease name and the effects of the mutations (*e.g*., silent, missense, insertions or deletions), *etc*.

### Interactive Drug Docking

VRmol performs drug docking through a built-in tool, AutoDock Vina [36]. In particular, VRmol implements interactive docking by specifying a restriction area defined by a 3D bounding box by users. Upon receiving the main structure, the drug structure and the restriction 3D bounding box coordinates, MGLTools (http://mgltools.scripps.edu/) is used for preprocessing the PDB files for docking. Then the built-in Vina program in the VRmol server starts to perform docking as a multi-thread task, which usually takes less than one minute to complete. The resulting docking structures are saved as individual PDB files by OpenBabel (http://openbabel.org/wiki/Main_Page), and can be loaded into the scene for visulization.

### Speech Recognition to Operate VRmol

Voice commands are converted into Base64-encoded characters and sent to the VRmol server by users’ web browsers. VRmol used the Baidu API (http://ai.baidu.com/) as the speech recognition engine for decoding, with fine-tuning on the frequently used words in structural biology studies, such as “*hydrogen*”, “*hydrophobicity*”, “*solvent excluded*”, *etc*., in the command list (Supplementary Table S4). Then, the decoded words are recognized as specific commands in VRmol by using an algorithm based on semantic similarity. In the semantic similarity searching procedure, the decoded words and all VRmol commands are uploaded onto an external knowledge base, WordNet [37], for calculating the similarity scores. The command with the highest similarity score is considered as the target command. The accuracy of speech recognition and command decoding has been remarkably improved upon addition of model training on the commands sets.

## RESULTS

VRmol uses cutting-edge WebVR technologies to model macromolecular structures in a virtual three-dimensional environment, making it to be a powerful tool for structural studies. It implements most commonly-used structural visualization styles and analysis functions as main stream desktop tools, with the supporting of cryo-EM density maps (Table 1). All of these analysis can be performed in two modes: the desktop mode and the VR mode. We provide a video for exploring the structure in VR mode (Video S1). As a cloud-based application, VRmol is seamlessly integrated with many online databases for structure-based disease and drug-related researches. A full description of the functions is available online at: https://VRmol.net/docs.

**Table 1.**
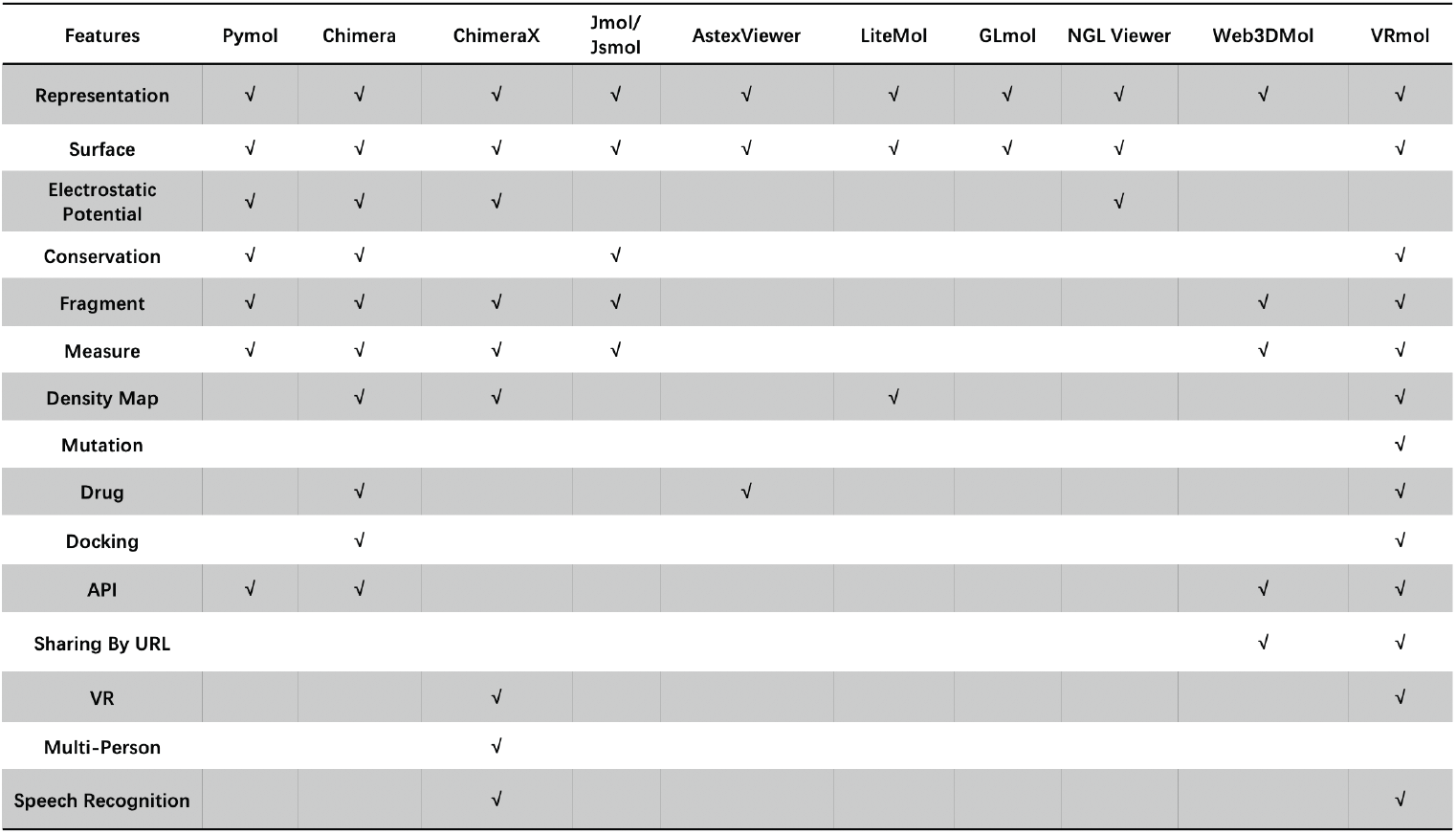
Features comparison bewteen VRmol and other structural visualization tools.

| Features | Pymol | Chimera | ChimeraX | Jmol/Jsmol | AstexViewer | LiteMol | GLmol | NGL Viewer | Web3DMol | VRmol |
| --- | --- | --- | --- | --- | --- | --- | --- | --- | --- | --- |
| Representation | √ | √ | √ | √ | √ | √ | √ | √ | √ | √ |
| Surface | √ | √ | √ | √ | √ | √ | √ | √ |  | √ |
| Electrostatic Potential | √ | √ | √ |  |  |  |  | √ |  |  |
| Conservation | √ | √ |  | √ |  |  |  |  |  | √ |
| Fragment | √ | √ | √ | √ |  |  |  |  | √ | √ |
| Measure | √ | √ | √ | √ |  |  |  |  | √ | √ |
| Density Map |  | √ | √ |  |  | √ |  |  |  | √ |
| Mutation |  |  |  |  |  |  |  |  |  | √ |
| Drug |  | √ |  |  | √ |  |  |  |  | √ |
| Docking |  | √ |  |  |  |  |  |  |  | √ |
| API | √ | √ |  |  |  |  |  |  | √ | √ |
| Sharing By URL |  |  |  |  |  |  |  |  | √ | √ |
| VR |  |  | √ |  |  |  |  |  |  | √ |
| Multi-Person |  |  | √ |  |  |  |  |  |  |  |
| Speech Recognition |  |  | √ |  |  |  |  |  |  | √ |

### Representation

VRmol supports various structural representation styles, including ribbon, tube, stick and ball & stick styles (Figure 2A), as well as structure surface, like solvent-accessible and Van de Waals surfaces (Figure 2B). Particularly, in the surface representation, users could variedly color the structural surface by a specific mode like by the hydrophobicity or secondary structure, and explore the meshed or transparent surface. A rich set of camera settings, lights, molecular surface material, rendering strategies, and model animations allow versatile structural visualization. Importantly, the minimal modeling units in VRmol are atoms, which enables high-resolution manipulation and editing of structures at the amino acid scale. In addition, cryo-EM density maps are well-supported and can be presented as different visualization styles like surface, slice, solid or mesh styles (Figure 2C), and the representation threshold can be adjusted by the control panel in user interface. In VRmol, users can interactively perform structural analysis using various operation options like selection, labeling, calculation and editing using VR menu in an immersive VR environment (Figure 2D, Supplementary Figure S1). To balance the requirement for structure modeling with high resolution and efficiency, VRmol automatically calculates a vision sphere, and only the objects within this sphere are modeled and rendered in high resolution.

**Figure 2.**
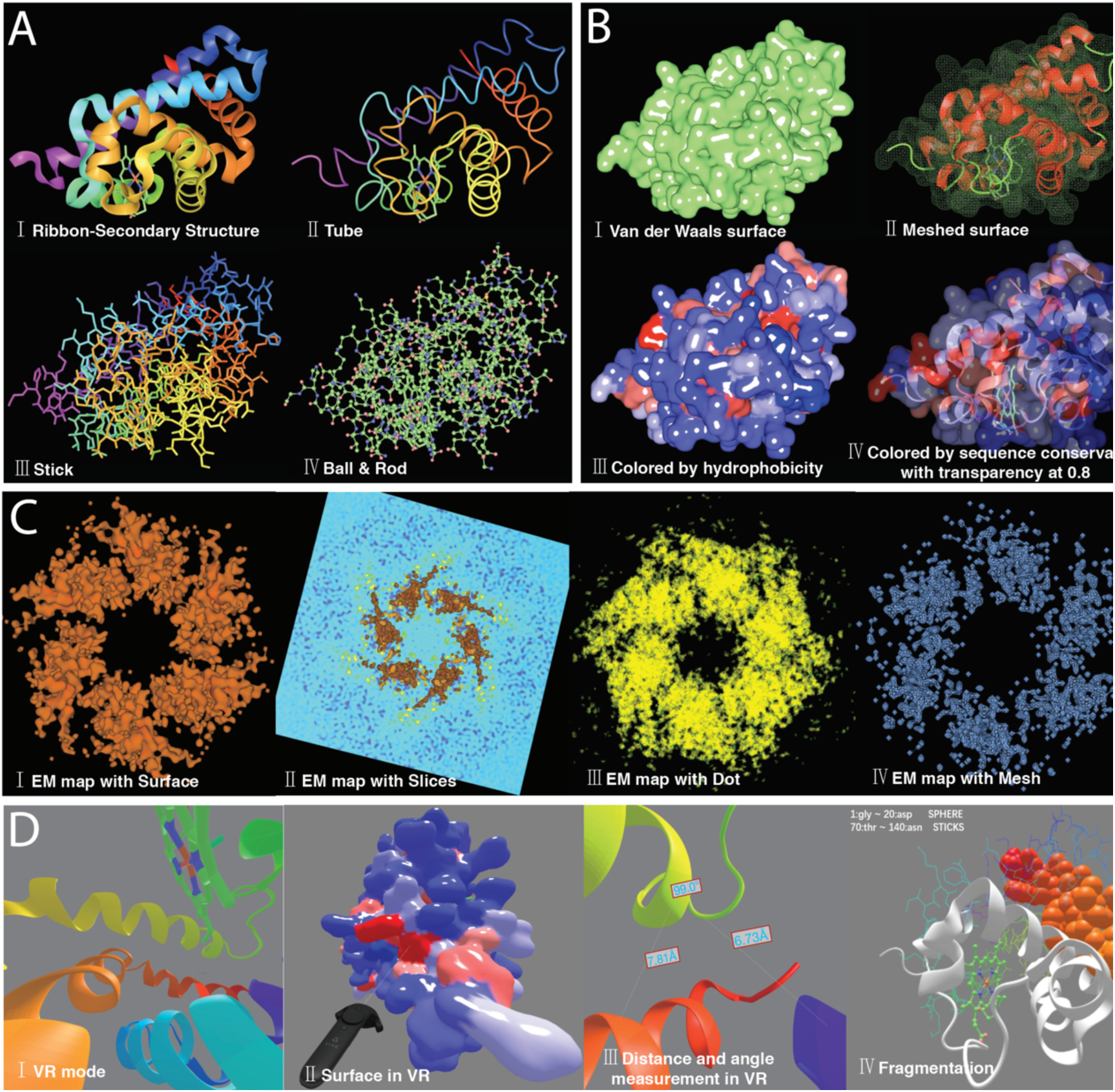
Structural representation in desktop and VR mode in VRmol. A. Seal myoglobin (PDB: 1MBS) [38] is presented with different representation styles including ribbon (I), tube (II), stick (III) and ball & stick colored by atom type (IV) in desktop mode. B. Seal myoglobin (PDB: 1MBS) is presented with different surface styles including Van der Waals surface (I), Meshed Van der Waals surface with secondary structure representation (II), Van der Waals surface colored by hydrophobicity (III) and Van der Waals surface colored by sequence conservation with transparency at 0.8 (IV) in desktop mode. C. The electron density map of human p97 (EMD: 3298, PDB: 5FTM) [39] is presented with four representation modes: surface (I), surface & slice (II), dot (III) and mesh (IV) in desktop mode. D. Representation styles of seal myoglobin (PDB: 1MBS) in VR mode, including ribbon (I), Van der Waals surface colored by its hydrophobicity(II), distance and angle measurement (III) and selected fragments variedly presented by Sphere and Sticks styles(IV).

### Structure Editing

Structure editing is one of the fundamental analysis methods in structural biology, especially in model building and mutation analysis. In VRmol, users could easily edit a structure by choosing the residue to be replaced and selecting the target amino acid type in the “Editing panel” (Figure 3A), and the edited amino acid could be highlighted by a distinct respresentation style. As shown in Figure 3B and C, the first amino acid, glycine, in chain a of seal myoglobin was replaced by threonine, and the resulting threonine was highlighted by being presented as sphere style in the edited structure. VRmol also provides a favorable function, called “*Fragment*”, facilitating to emphasize some specific regions by presenting them in diverse styles when perform structural visualization and analysis.

**Figure 3.**
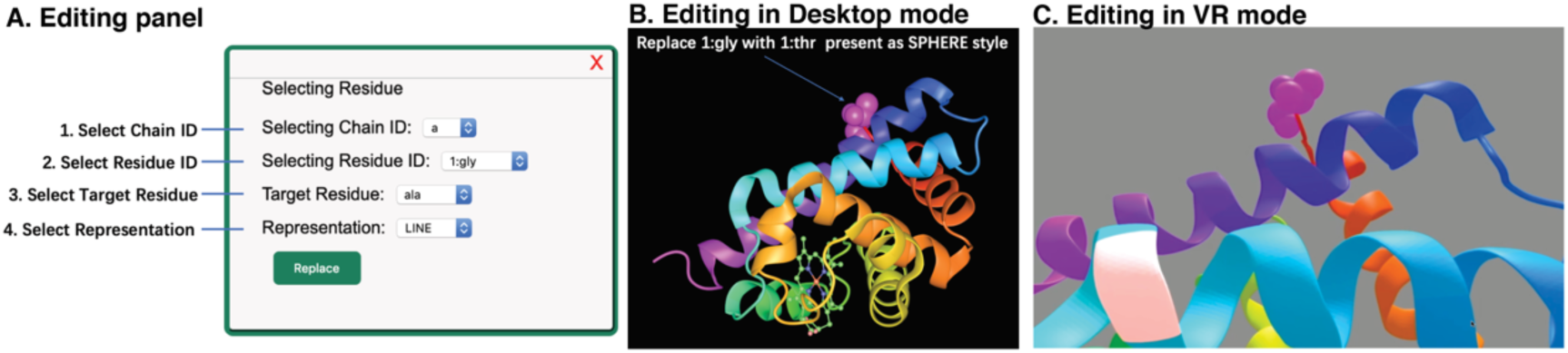
Structural editing in VRmol. A. Editing panel in Desktop mode, where users could select the original residue and the target residue type to perform editing. B. Editing results in desktop mode. The first residue (glycine) in Seal myoglobin (PDB: 1MBS) was replaced by threonine. C. Editing results in VR mode.

### Visualization of Genomic Variations

Genomic variations could alter protein folding, leading to aberrant functions. Analyzing the distribution of genomic variations on protein structures can shed light on the genetic basis of many complex diseases. VRmol facilitates translation research by automatically retrieving data from a list of genomic variation databases, including TCGA, CCLE, ExAc, and dbSNP, and mapping mutations onto structures (Figure 4A). This approach can reveal whether and where frequently mutated residues spatially cluster on a structure. As an example, Figure 4B shows genomic variation sites on the structure of human glucose transporter GLUT1 (PDB id: 4PYP) [40], highlighted by red balls. In the “Mutation info table” in the user interface, there listed the mutated amino acid numbers and types, variation related diseases and variation results, like silence or missense variations. We can conclude that genomic variations have been reported to happen in A354, S365 and Y366 of GLUT1, and such variations lead to different phynotypes or diseases. In the VR mode, users could also clearly visualize these variation sites on structure model labeled by colored balls (Figure 4C).

**Figure 4.**
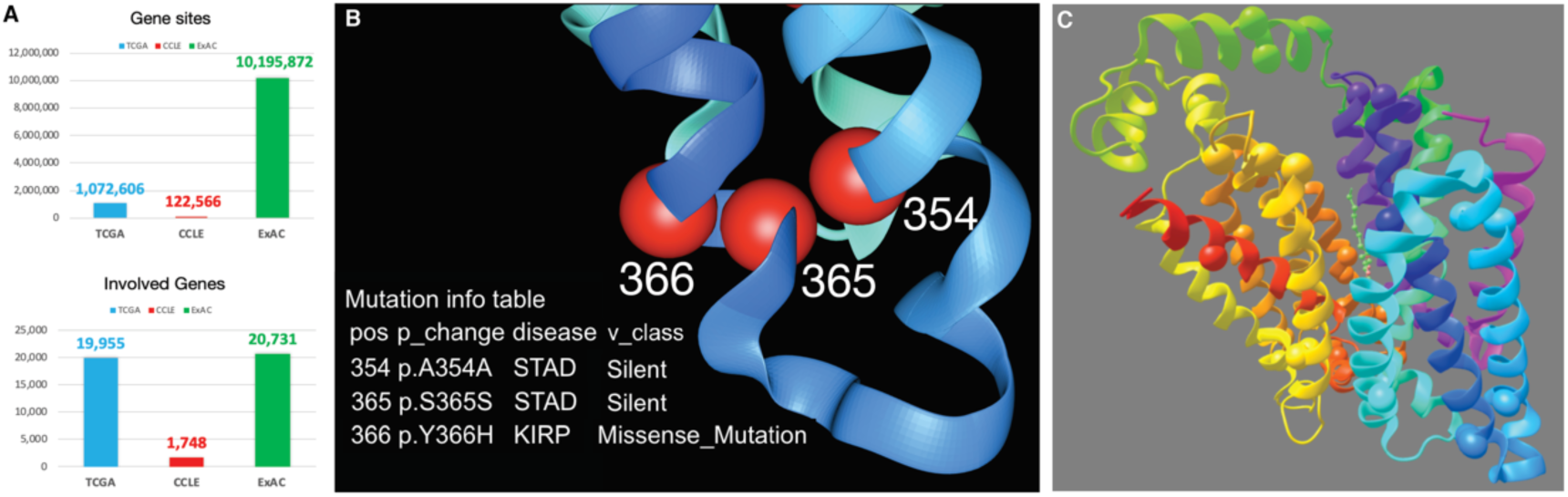
Genomic variations visualization in VRmol. A. Statistics of variation sites and involved genes in three genomic variation databases (TCGA, CCLE, ExAC). B. Three variation sites (from TCGA) mapped onto the structure of human glucose transporter GLUT1 (PDB id: 4PYP) were highlighted by red balls with detailed mutation information listed aside in desktop mode. C. Genomic variations sites (from TCGA) were mapped onto the structure of GLUT1 and highlighted by colored balls in VR mode.

### Interactive Drug Docking

The VR environment offers an opportunity for intuition-guided docking of drugs or other drug-like small molecules onto target structures. VRmol can automatically search and load relevant drug molecules or small ligands of the target structure from multiple databases, such as DrugBank, ChEMBL, BindingDB [41], SwissLipids [42], and GuidetoPHARMACOLOGY [43] (Supplementary Table S3). VRmol can perform global drug docking in the whole space of a target structure (Figure 5A). Alternatively, users can do guided-docking by entering the “drug dragging” mode to move a molecule to potential binding sites on the target structure and then manually select surrounding areas (generating 3D box coordinates) as drug searching regions (Figure 5B). VRmol uploads the structure files and the box coordinates to the cloud-based VRmol server for automatic searching and docking. The nine best docking modes are returned with docking scores. Normally, the docking mode with lower score is more reliable and meaningful. As an example, Figure 5C shows a drug molecule, N-Formylmethionine (DrugBank id: DB04464), docking in a pocket of linear di-ubiquitin (PDB id: 2W9N) [44]. Video S2 shows the operation for docking in VR step by step. The di-ubiquitin was rendered by hydrophobicity surface, while the drug molecule by ball&still style. Nine docking results were listed on the right side, and by selecting the top one with lowest score, we could find that N-Formylmethionine was finely docked in the target pocket.

**Figure 5.**
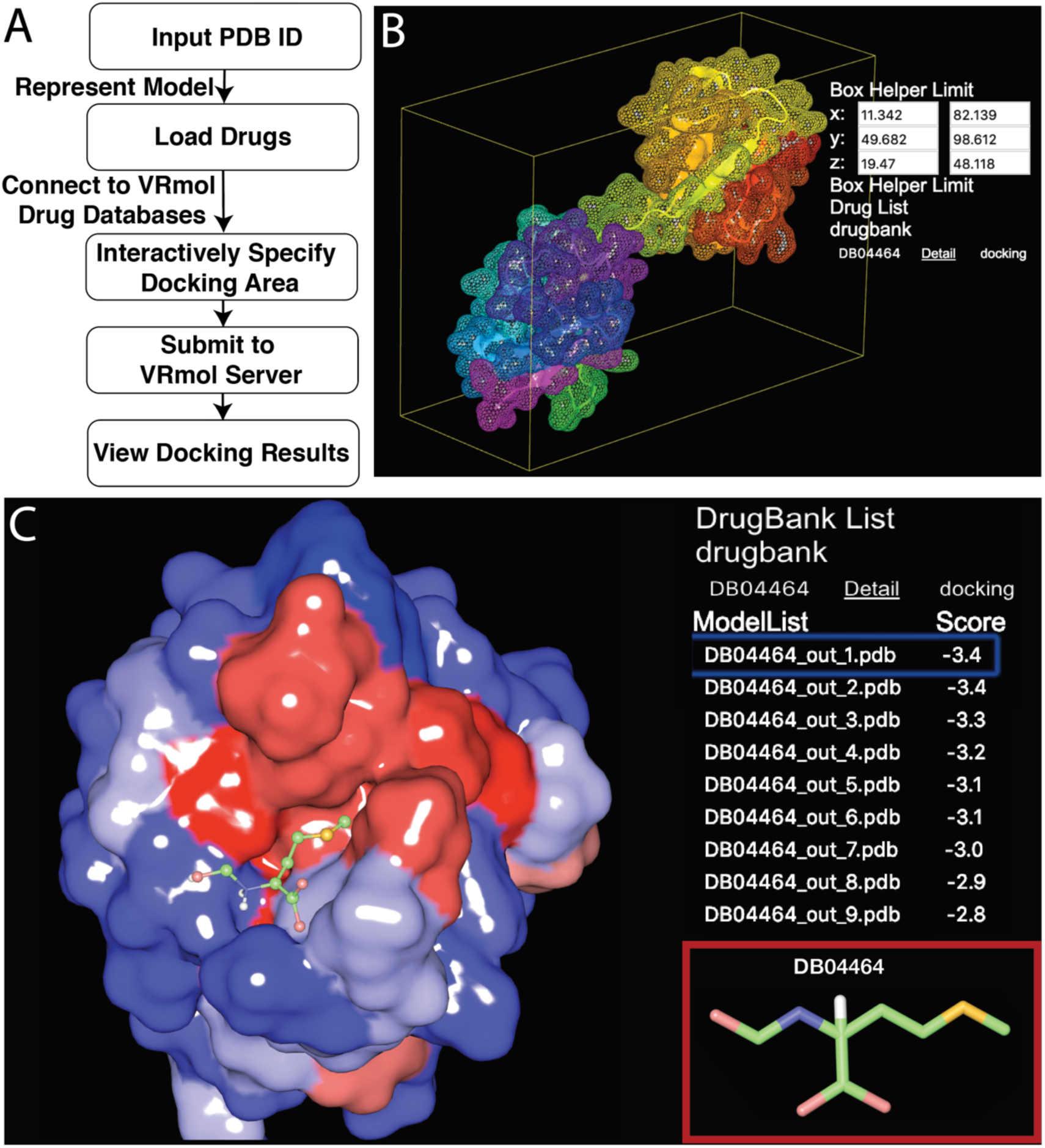
Drug Docking in VRmol. A. The pipeline of interactive docking in VRmol. B. Interactively specify a restriction area. C. Docking of the drug DB04464 from DrugBank onto di-ubiquitin (PDB id: 2W9N). Nine potential docking modes were listed on the right.

### Speech Control and Easy Collaboration

For convenient interaction, VRmol implemented a speech recognition function with over 50 spoken operation commands in English that cover most functionalities (Supplementary Table S4). Operating by voice commands makes VRmol to be a friendly and practical tool to explore structural world not only in structural biology research but in teaching or scientific popularization. Indeed, VRmol included both English and Chinese options for speech control in VR mode (Details in https://vrmol.net/docs/#header-n274). For scientific collaboration in structural biology field, reseachers tend to share specific views of an individual structure on a visualization scene with their collaborators. However, scene sharing in desktop-based tools, such as Pymol and Chimera, is usually achieved by saving scenes as 3D geometry files with a large size and with monotonous rendering, thus making it inconvenient or inappropriate to share scenes. In contrast, VRmol, as a cloud-based application, can share a VR scene via URL, thereby improving collaborative communications. VRmol encodes and renders individual atoms separately, with settings and styles being able to be retrieved and imported into the scene when the URL is remotely loaded. Users can specify and alter the structural representation styles in real time when sharing scenes, by choosing the corresponding options in the URL.

## DISCUSSION

The VRmol platform is an integrative cloud-based system that uses VR technology to visualize and analyze macromolecular structures. Users can explore molecular structures in a fully immersive 3D virtual environment by using a VR-enabled device, without the need to install other packages. VRmol harbors many new functions that do not exist in other VR-supported structural biology tools. These features include various structural visualization and analysis functions (e.g., structure editing), rich interactive methods (e.g., speech recognition), and easy collaboration accomplished by URL sharing. Importantly, VRmol connects to many disease-related mutation databases and drug databases, and thus provides an integrative platform for translational researches. Collectively, we envision that VRmol can largely benefit not only structural biology research but also scientific popularization, and has extensive potential applications on structure-related diseases study and corresponding drugs discovery.

## DATA AVAILABILITY

VRmol is freely available at https://VRmol.net, and its tutorial at https://vrmol.net/docs/. VRmol source code is available at http://github.com/barrykui/VRmol.

## SUPPLEMENTARY DATA

Supplementary Data are available at NAR Online.

## AUTHOR CONTRIBUTIONS

Q.C.Z. and H.W. conceived this project. Q.C.Z. supervised the project. K.X. implemented VRmol with the help from J.X., N.L. and C.G. tested VRmol. K.X., N.L., H.W., and Q.C.Z. wrote the manuscript with inputs from all authors.

## ACKNOWLEDGMENTS

We thank Haitao Li (Tsinghua University) for helpful comments on the development of VRmol. We also thank Maoxiang Shi for useful suggestions to improve the efficiency of VRmol.

## Supporting Information

**Video S1. Three minutes video for exploring the structure in VR mode.**

**Link: https://vrmol.net/docs/v/vr/overview.mp4**

**Video S2. Four minutes video for docking drug into the structure in VR mode.**

**Link: https://vrmol.net/docs/v/vr/docking.mp4**

**Figure S1.**
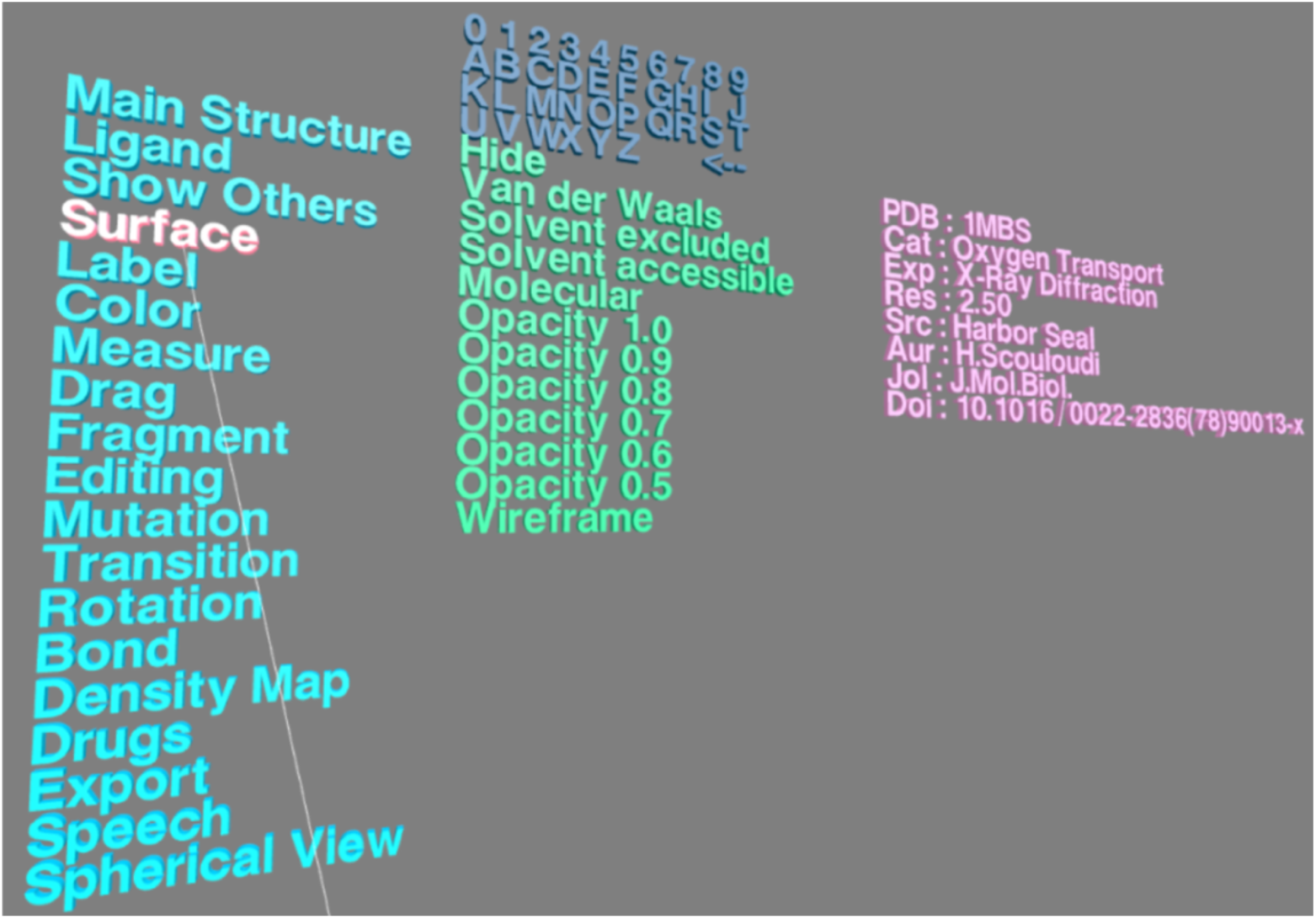
VR menu in the VR mode of VRmol. The VR Menu contains three parts. Virtual keyboard composed of 26 alphabet, 10 numbers, an execution button and a deletion button is on the top. On the bottom are a menu and an information panel, respectively.

**Figure S2.**
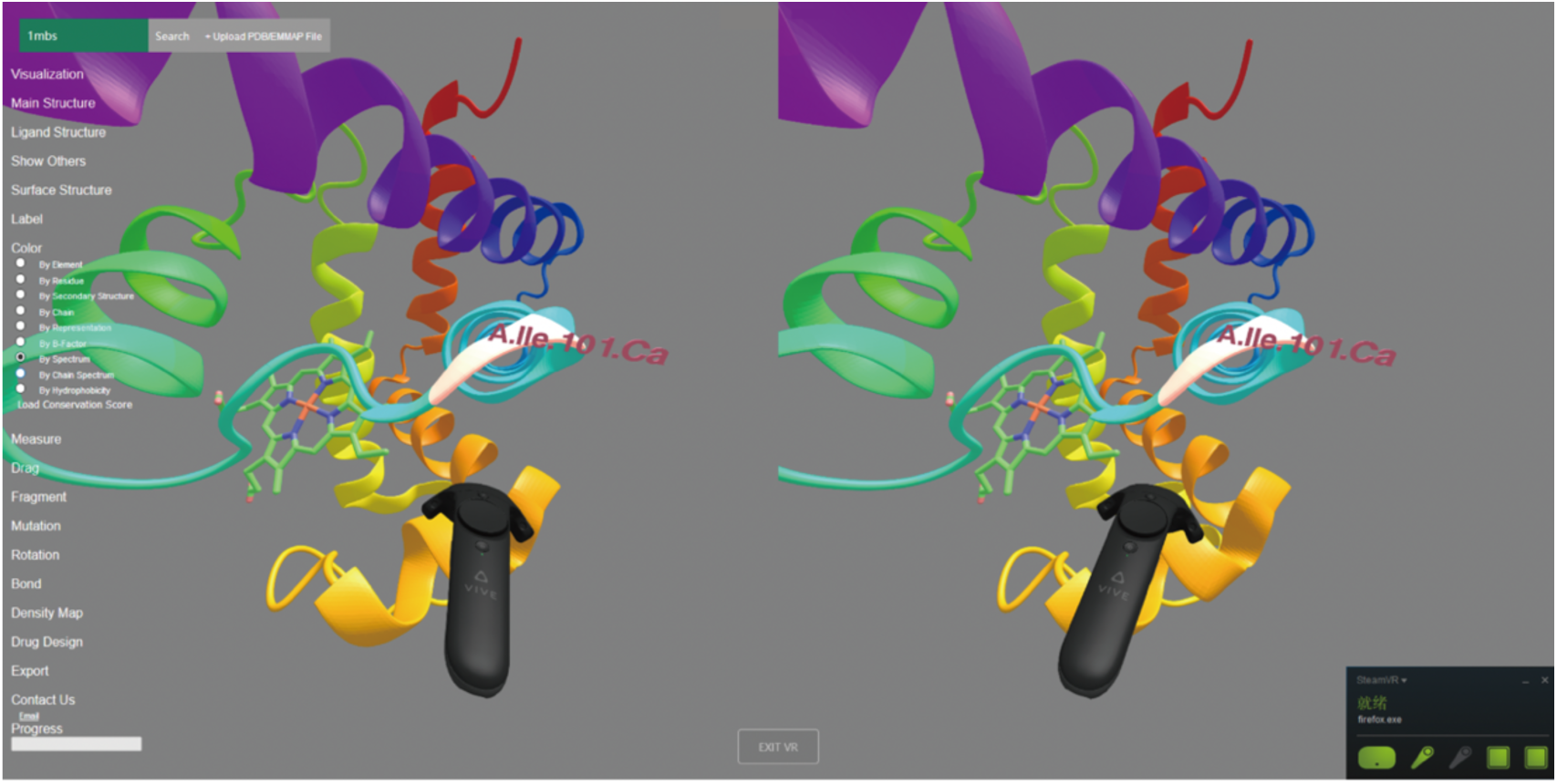
Screenshot of the stereo view in VR mode.

**Figure S3.**
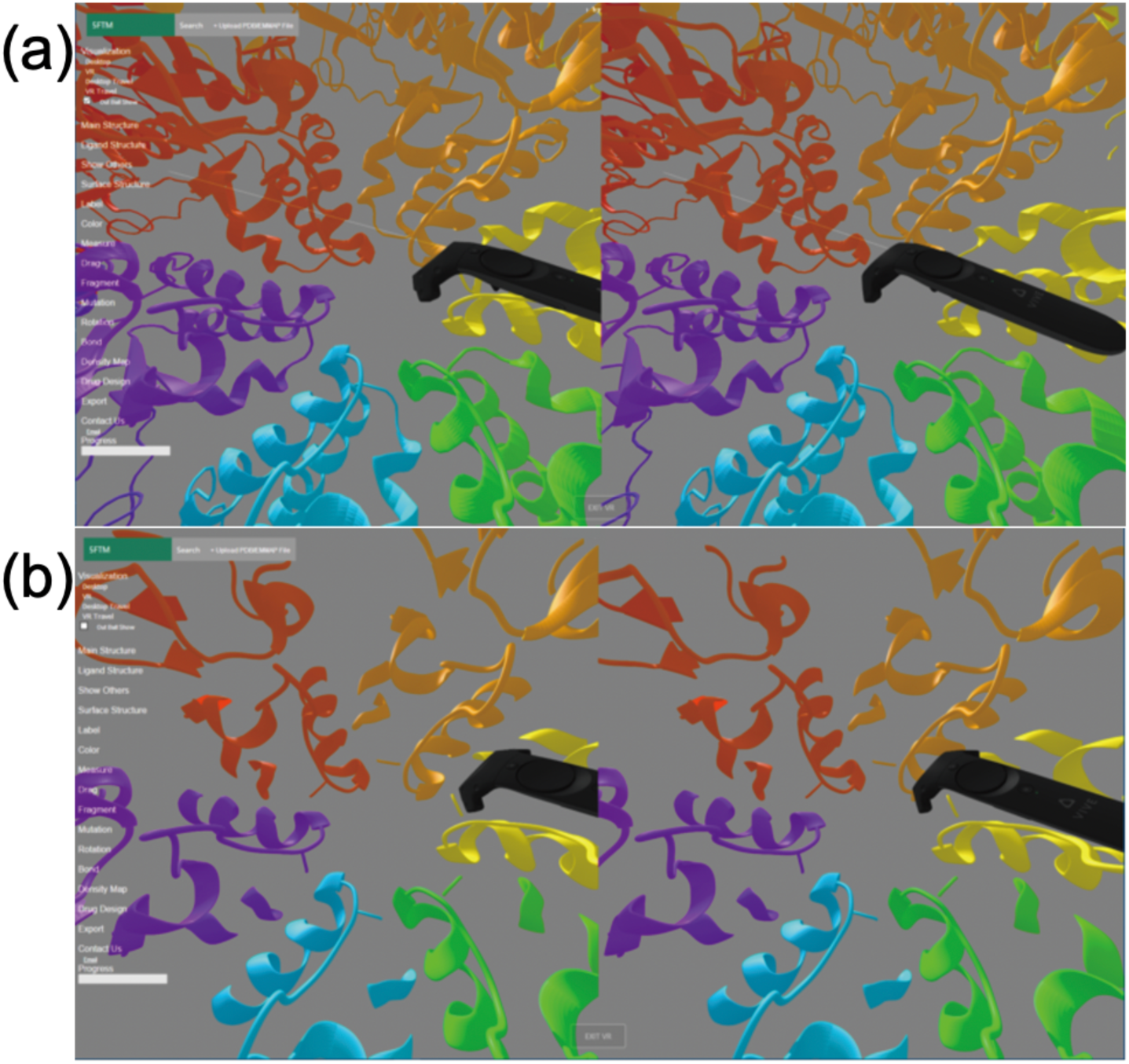
Spherical View. (a). Ribbonlike structures outside of the spherical view is shown with low resolution. (b). Ribbonlike structures outside of the spherical view are hided.

**Table S1:** Key resources applied in VRmol.

| REAGENT or RESOURCE | SOURCE | IDENTIFIER |
| --- | --- | --- |
| **Deposited Data** |  |  |
| TCGA | [1] | https://portal.gdc.cancer.gov/ |
| ExAC | [2] | http://exac.broadinstitute.org/ |
| CCLE | [3] | https://portals.broadinstitute.org/ccle/about |
| dbSNP v150 | [4] | https://www.ncbi.nlm.nih.gov/projects/SNP |
| DrugBank | [5] | https://www.drugbank.ca/ |
| ChEMBL | [6] | https://www.ebi.ac.uk/chembl/ |
| BindingDB | [7] | https://www.bindingdb.org/bind/index.jsp |
| SwissLipids | [8] | http://www.swisslipids.org/ |
| GuidetoPHARMACOLOGY | [9] | http://www.guidetopharmacology.org |
| EMDB | [10] | https://www.ebi.ac.uk/pdbe/emdb/ |
| **Software and Algorithms** |  |  |
| Marching cube | [11] | http://webglmol.osdn.jp/surface.html |
| w3m.js | [12] | https://web3dmol.net/ |
| Three.js v95 | threejs.org | http://github.com/three.js |
| ConSurf | [13] | http://consurf.tau.ac.il/2016/ |
| AltPDB | AltPDB | https://github.com/AltPDB/protein-viewer |
| AutoDock vina | [14] | http://vina.scripps.edu/ |
| MGLTools | MGLTools | http://mgltools.scripps.edu/ |
| openbabel | OpenBabel | http://openbabel.org/wiki/Main_Page |

**Table S2.** VR devices supported in VRmol.

| Device | Frame Rate<br>(frame per second) | OS | Web browser |
| --- | --- | --- | --- |
| HTC VIVE | 90 | Windows 7 + | Firefox Nightly |
|  |  | macOS | Firefox Nightly |
| Windows Mixed Reality | 90 | Windows 10<br>(build 1709+) | Microsoft Edge |
|  |  | Windows 10 | Firefox |
| Oculus Rift | 90 | Windows 10 | Firefox |
| Google<br>Cardboard | <60 | Android | Chrome |
|  |  | iOS | Safari |
| Google<br>Daydream | <60 | Android | Chrome |
| Mojing | <60 | Android | Chrome |

**Table S3.** Drug and small molecule ligand databases supported in VRmol.

| Database | Count of the targets/drugs | Latest Release |
| --- | --- | --- |
| ChEMBL | 11,538 / 2,101,843 | May 2017 |
| DrugBank | 4,096 / 9,588 | 06 July 2017 |
| BindingDB | 7,225 / 621,060 | 01 September 2017 |
| SwissLipids | 506,497 / 7,835 | 13 September 2017 |
| GuidetoPHARMACOLOGY | 1,684 / 1,334 | 22 August 2017 |

**Table S4.**
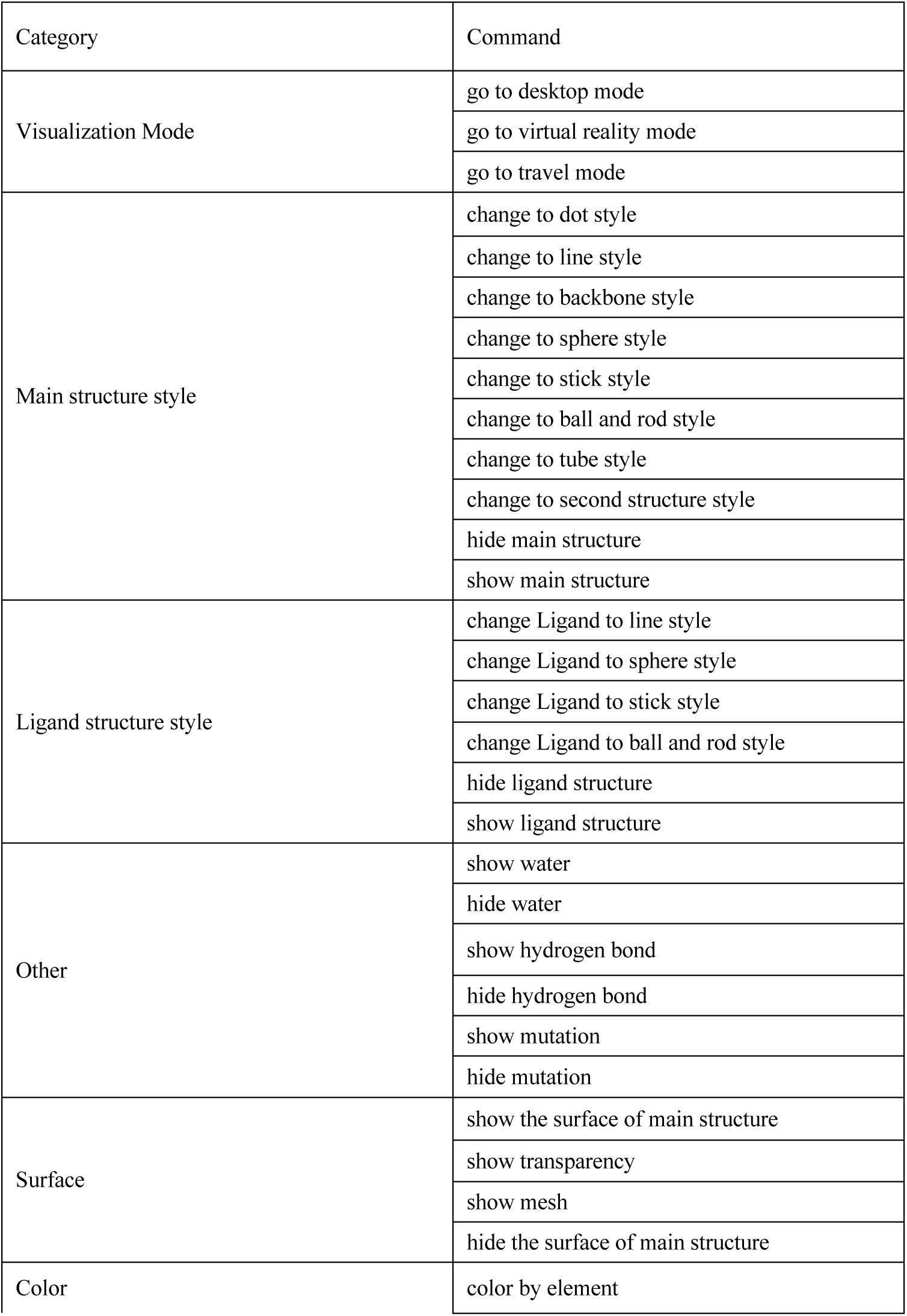
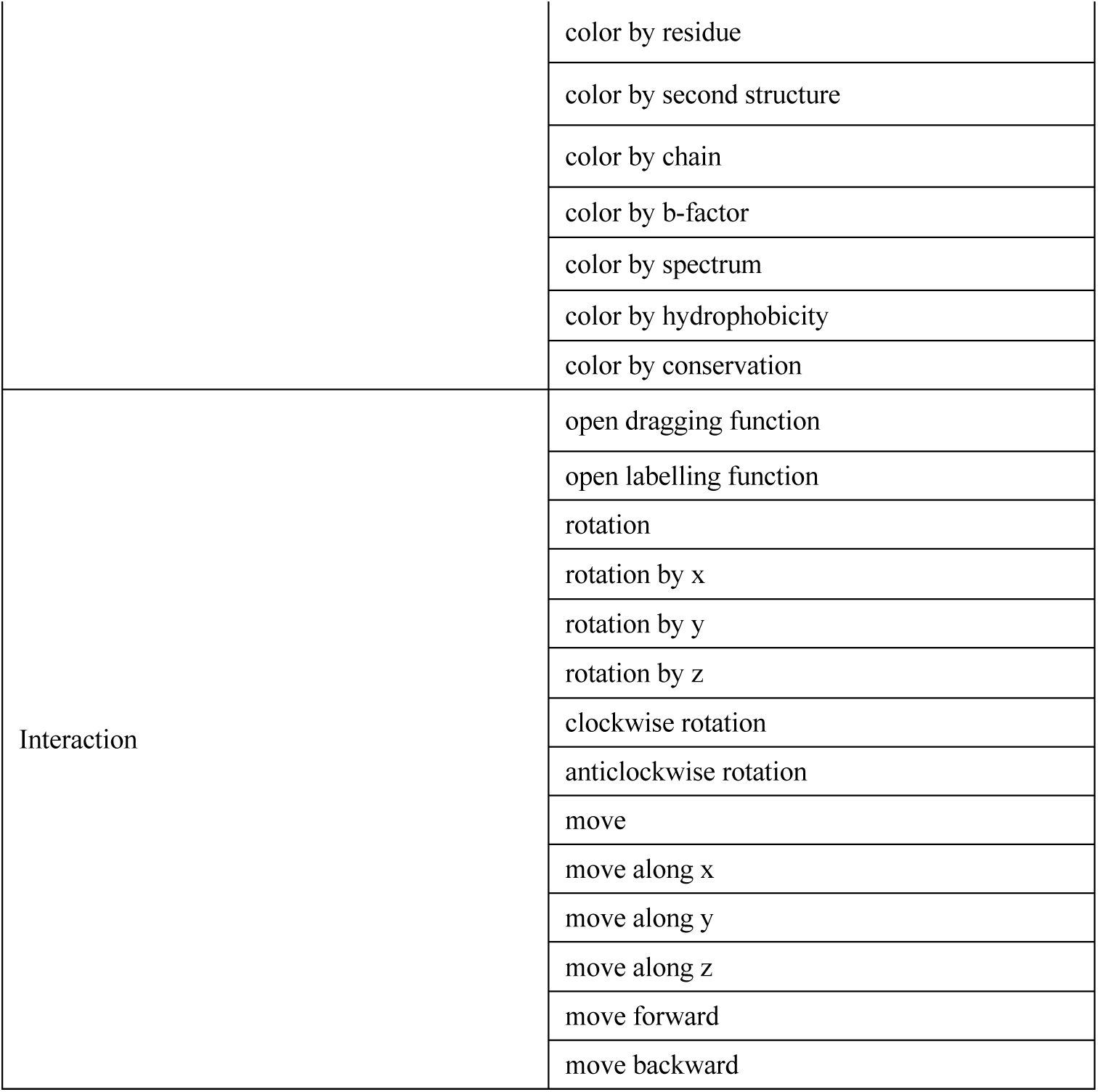
Commands in speech recognition.

